# Detecting changes in the *Caenorhabditis elegans* intestinal environment using an engineered bacterial biosensor

**DOI:** 10.1101/717215

**Authors:** Jack W. Rutter, Tanel Ozdemir, Leonor M. Quintaneiro, Geraint Thomas, Filipe Cabreiro, Chris P. Barnes

## Abstract

*Caenorhabditis elegans* has become a key model organism within biology. In particular, the transparent gut, rapid growing time and ability to create a defined gut microbiota make it an ideal candidate organism for understanding and engineering the host microbiota. Here we present the development of an experimental model which can be used to characterise whole-cell bacterial biosensors *in vivo*. A dual-plasmid sensor system responding to isopropyl *β*-D-1-thiogalactopyranoside was developed and fully characterised *in vitro*. Subsequently, we show the sensor was capable of detecting and reporting on changes in the intestinal environment of *C. elegans* after introducing exogenous inducer into the environment. The protocols presented here may be used for aiding the rational design of engineered bacterial circuits, primarily for diagnostic applications. In addition, the model system may serve to reduce the use of current animal models and aid in the exploration of complex questions within general nematode and host-microbe biology.

## Introduction

As synthetic biology is increasingly applied to develop microbiome engineering tools such as biosensors [1] and live biotherapeutics [2, 3], animal models will be vital for building systems that function robustly in complex *in vivo* environments. As such, there is a demand for cheap, robust, tractable model systems that can serve both as a development platform and also to probe host-microbe interactions.

The mouse is currently the most widely used model to study the intestinal microbiome. The similar taxonomic levels of the microbiota to humans, an extensive knowledge of genetic backgrounds, custom genotypes and phenotypes, plus the use of humanised gnotobiotic systems all lead to a mimicry of the human gut microbiota phylogenic composition and allow researchers to investigate perturbations in a human-like system [4, 5, 6, 7]. To date, the majority of synthetic biology approaches to engineer or monitor the intestinal microbiota *in vivo* have been demonstrated within a mouse model [8, 9, 10, 11, 12, 13, 14]. However, the majority of the cross-talk between the gut microbiota and the host is host-specific [15]. Furthermore, there is a problem with non-reproducible findings due to genetic variability, handling techniques, mouse vendors and diet [16, 17].

The invertebrate *Caenorhabditis elegans* is a transparent nematode worm 1 mm in length that lives in temperate soil environments and feeds on soil bacteria. It was the first multicellular organism to have its whole genome sequenced [18] and a combination of a short 2-3 week lifespan, transparent cell wall and genetic tractability have enabled it to become an extremely versatile model system used to study energy metabolism, immunity and ageing [19]. While the *C. elegans* worm demonstrates a diverse microbiota in the wild [20, 21], it is typically monoxenically grown with one species in the lab; this enables researchers to easily create and maintain a defined intestinal microbiota. The intestines are one of their major organs and constitute roughly a third of their somatic mass [22]. The transparent cell wall and aerobic lumen also enable the simple visualisation of fluorescent proteins and markers. The worm possesses an innate immune system which is used to regulate the intestinal bacterial load as it ages [23]. Peak transition of bacteria through the intestines can be as short as 2 minutes during young adulthood, although the bacteria eventually colonise the lumen in a number of days as the worm ages [24]. The emerging need for both convenient and robust tools to investigate host-microbiota interactions has resulted in a growing interest in the use of *C. elegans* as a live animal model for both host-microbiota interactions and synthetic biology. Examples include high-throughput screens to elucidate the complexity underlying host-microbe-drug interactions [25], understanding how bacterial produced metabolites affect worm gene expression and its lifespan [26], and the role of stochasticity in the colonisation of the gut by microbiota [27]. In synthetic biology, *C. elegans* has been used as a target for engineered nematicidal bacteria [28], and to characterise a sense-and-kill synthetic circuit in *E. coli* Nissle 1917 (EcN) that could colonise the worm to prevent a *Pseudomonas aeruginosa* gut infection [29].

Here we describe the development of protocols for the characterisation of bacterial whole-cell biosensors that function within the *C. elegans* gastrointestinal tract. We first constructed a dual plasmid-based biosensor system in EcN that responds to isopropyl *β*-D-1-thiogalactopyranoside (IPTG). This biosensor was then characterised *in vitro* at 37°C and 21°C. We then describe experimental protocols and an automated imaging pipeline that allowed us to examine quantitatively how this biosensor strain reports on changes in environmental levels of IPTG. Taken together these results outline how *C. elegans* can be used as a novel model organism for further characterising biosensors and host-microbe interactions within the digestive tract.

## Results and discussion

### Ratiometric, dual-plasmid IPTG biosensor system

A dual reporter system using mCherry and GFP was created in EcN (Figure 1A) with mCherry constitutively expressed and GFP expression under the control of the inducible pLac promoter, allowing for the calculation of a ratiometric fold increase in GFP induction (plasmid maps are given in Figure S1). This approach is considered to be more effective in reducing background noise and accounting for fluctuations in diverse conditions *in vivo* [30, 31]. The dual mCherry and GFP system was initially characterised within EcN, in liquid LB culture at 37°C (preliminary characterisation in LB and M9 given in Figures S2 and S3). In comparison to the negative promoterless EcN_OG241_GFP_mCherry and positive constitutive EcN_OXB19_GFP_mCherry controls, where the GFP:mCherry ratio was seen to remain relatively constant (Figure S4), the pLac inducible system showed a robust increase in the GFP:mCherry ratio upon induction of the dual reporter system (Figure 1B and 1C).

**Figure 1:**
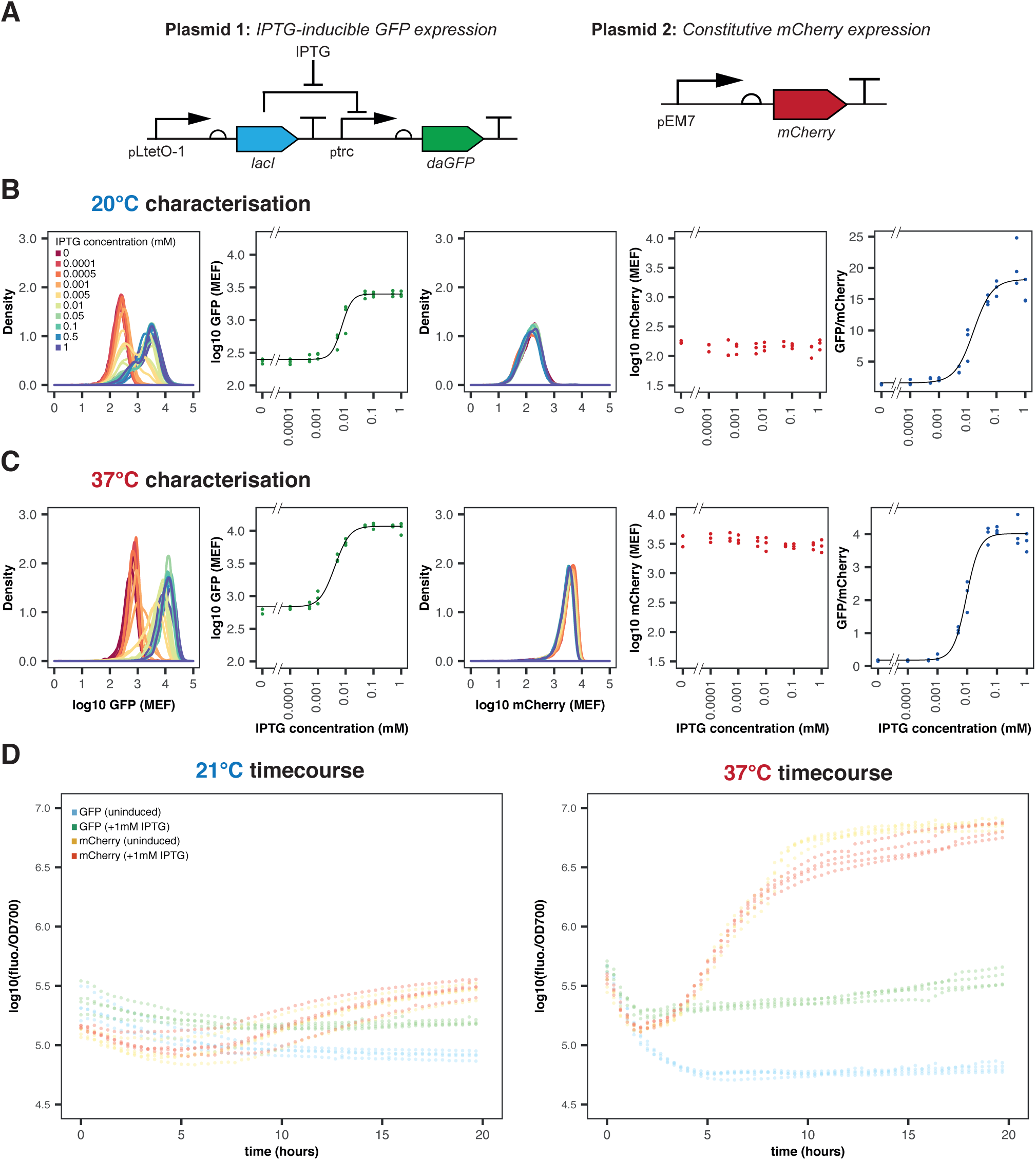
EcN_pLac_GFP_mCherry *in vitro* characterisation, within LB media at 20°C and 37°C. (**A**) Plasmids in the biosensor system, based on constitutive mCherry and inducible GFP expression. (**B**) and (**C**) characterisation of the strain at 20°C and 37°C, respectively. From left to right: density plot of GFP induction, median GFP fluorescence, density plot of mCherry fluorescence, median mCherry fluorescence and GFP:mCherry ratios over all IPTG concentrations. Flow cytometry data with 10 000 events (n=3). (**D**) Timecourses of both uninduced and induced GFP (blue/green) and mCherry (orange/red), respectively, induced at 3 hours growth. (n=4, circles indicate individual datapoints).

In order to try and mimic conditions of the *C. elegans* digestive tract *in vitro*, characterisation was additionally performed at 21°C (room temperature). At 37°C, the median GFP fluorescence was found to increase from 684.57 ± 253.99 MEF (molecules of equivalent fluorophore, fitted value ± standard error, for 30,000 events from 3 biological replicates) when uninduced to 11721 ± 269.40 MEF after induction with 1 mM IPTG. A change in GFP expression was also detected at 20°C, increasing from 250.10 ± 60.93 MEF to 2505.53 ± 64.64 MEF with 1 mM IPTG induction. The median GFP:mCherry ratio for induction with 1mM IPTG was much greater at 20°C than at 37°C, 18.16 ± 0.84 and 4.01 ± 0.08, respectively. The threshold for detection, *K*_*d*_ and dynamic range (illustrated in Figure S5) of the EcN_pLac_GFP_mCherry strain were adversely affected at 20°C (Table 1). This was particularly prominent for the dynamic range of the GFP:mCherry ratios, which was approximately halved at 21°C when compared to 37°C (10.38 and 21.08, respectively). This may be because of slower growth and expression rates at the lower temperature. However, the linear range of the GFP:mCherry ratio was much larger at 21°C than 37°C; at 21°C the linear range spanned 72.86 *µ*M, from 0 −72.86 *µ*M; at 37°C the linear range only covered 35.65 *µ*M, from 0.16 −35.81 *µ*M. This suggests that the ratiometric output can be used to distinguish between a larger range of inducer concentrations at 20°C than 37°C.

**Table 1:** Hill parameter fitting to GFP induction and GFP:mCherry ratio curves, for both 20°C and 37°C

| Parameter | GFP induction curves |  | GFP:mCherry ratios |  |
| --- | --- | --- | --- | --- |
|  | 20°C | 37°C | 20°C | 37°C |
| $f^{min}$ (MEF) | 250.10±60.93 | 684.57±253.99 | 1.60±0.60 | 0.18±0.07 |
| $f^{max}$ (MEF) | 2505.53±64.64 | 11721.49±269.40 | 18.16±0.84 | 4.01±0.08 |
| $K_d$ (μM) | 11.17±1.17 | 8.68±0.76 | 15.93±3.47 | 9.39±0.70 |
| $n$ | 2.61±0.83 | 1.88±0.38 | 1.35±0.30 | 1.98±0.35 |
| dynamic range | 9.02 | 16.12 | 10.38 | 21.08 |
| linear range (μM) | 34.67 | 34.19 | 72.86 | 35.65 |

<sup>a</sup> (fitted values ± standard error where appropriate, given to 2 d.p.).

In summary, while there appear to be differences seen in biosensor performance between the two temperatures, the characterisation at 20°C showed that the EcN_pLac_GFP_mCherry biosensor was capable of detecting changes in IPTG concentration in conditions more commensurate with the temperature within the *C. elegans* digestive tract. Therefore, the biosensor was compatible with the handling and culture of the *C. elegans* nematode.

### Imaging of the colonised *C. elegans* intestinal environment

In order to test the hypothesis that *C. elegans* can be used to characterise bacterial biosensors, we developed the experimental protocol depicted within Figure 2A. Preliminary experiments confirmed that the wild type lab N2 *C. elegans* strain could indeed grow and develop successfully on both the EcN_OG241_GFP_mCherry and EcN_OXB19_GFP_mCherry control strains constitutively expressing mCherry or GFP and mCherry, respectively. In addition, the GFP signal could be used to image the *C. elegans* intestine. The negative GFP control EcN_OG241_GFP_mCherry showed only high levels of mCherry (Figure 2B). EcN_OXB19_GFP_mCherry showed direct co-localisation with strong expression of both GFP and mCherry throughout the digestive tract (Figure 2C).

**Figure 2:**
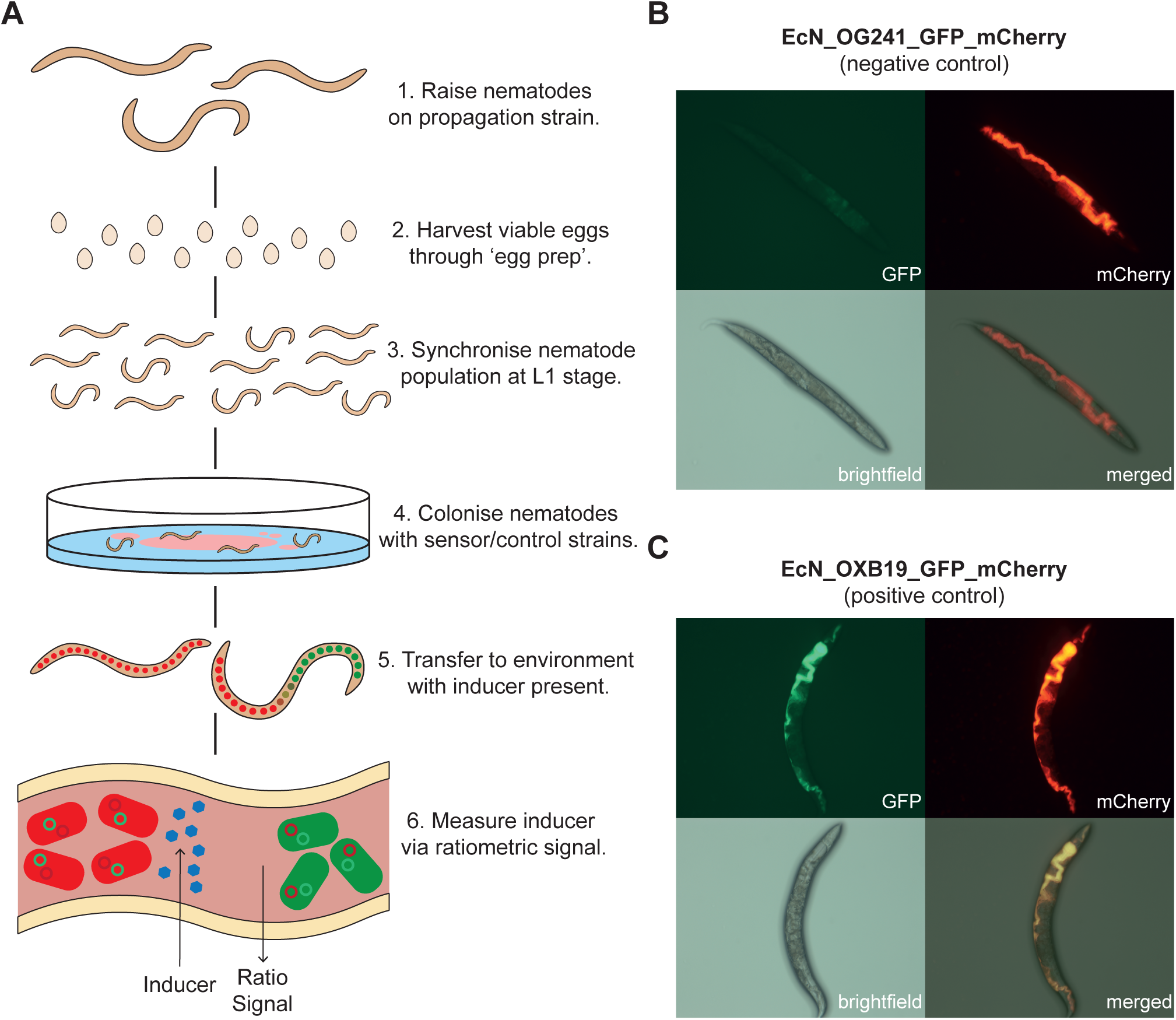
(**A**) Graphical representation of *C. elegans* induction assay protocol with EcN-NGM plates and sensor induction quantification using the GFP:mCherry ratio. Representative images of nematodes colonised with, (**B**) EcN_OG241_GFP_mCherry (negative control strain) expressing only mCherry and (**C**) EcN_OXB19_GFP_mCherry (positive control strain) constitutively expressing both mCherry and GFP. Panel labels refer to the imaging method.

Once it was clear that the nematodes could survive on the engineered EcN strains and that the reporter proteins could be detected, an image processing pipeline was developed that could be used to quantify the GFP:mCherry ratios (Figure 3A). The pipeline, developed in MATLAB, was able to successfully extract worm bodies from the images (ignoring noise from the plate background) and extract the colonised intestines of the worm images. Furthermore, steps were added which allowed for the discarding of uncolonised worm images (using a threshold on minimum mCherry fluorescence) and removal of GFP autofluorescence (subtraction of mean nematode body GFP intensity). It should be noted that for images which contained more than one nematode, the GFP:mCherry ratio was calculated over the entirety of the image. In addition, it was seen that not all nematodes became colonised and therefore some images were discarded during automated analysis (a representative image of an uncolonised nematode can be seen in Figure S6; overall, approximately 17% of the worm images collected for Figure 4 were discarded and deemed ‘uncolonised’ based on our mCherry threshold). This pipeline was initially trialled on nematodes which were grown on plates of EcN_pLac_GFP_mCherry +/− 1 mM IPTG, for 7 days before imaging (representative images can be seen in Figure 3B); the results of this experiment are presented in Figure 3C.

**Figure 3:**
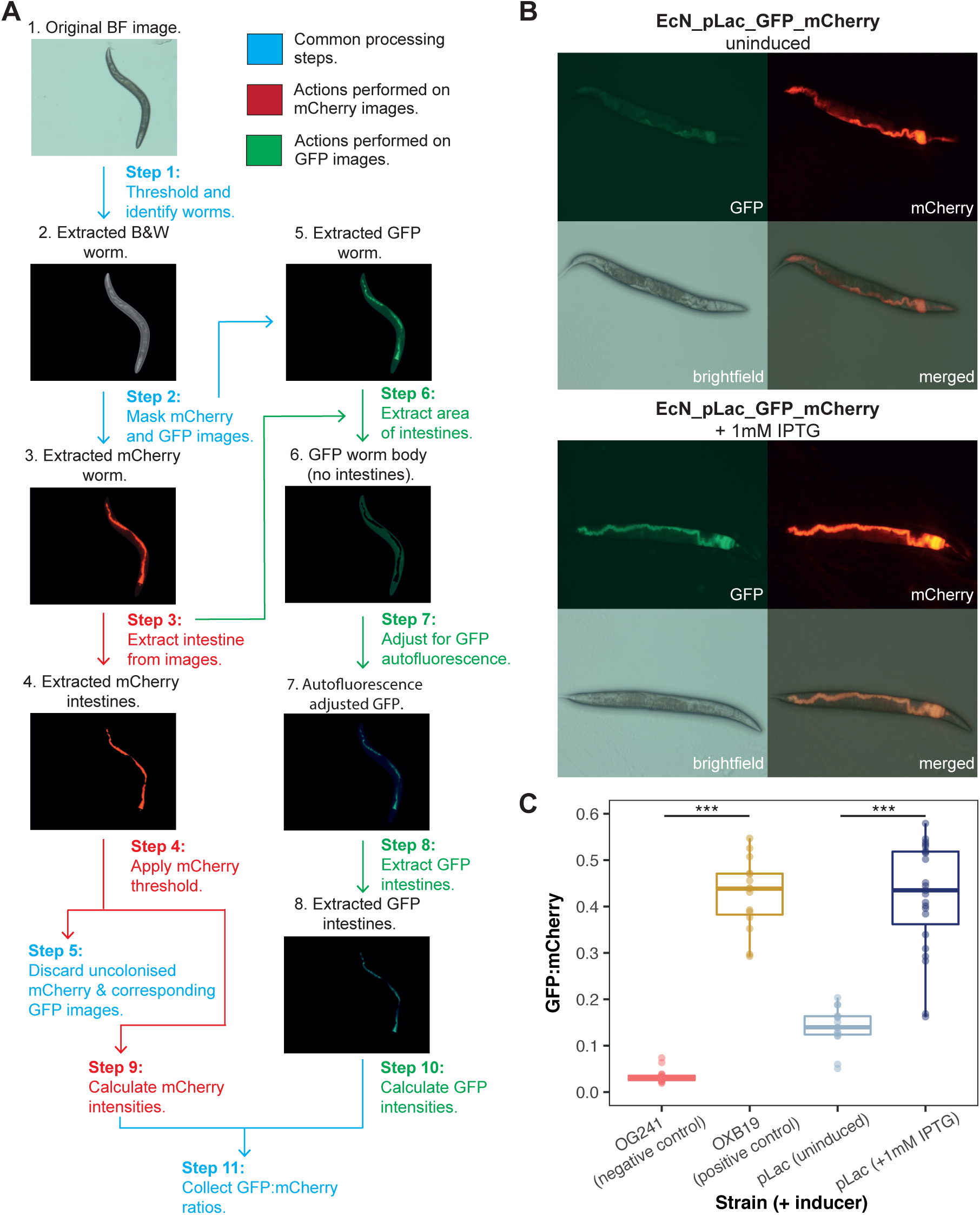
(**A**) Image analysis pipeline developed to automate quantification of biosensor induction (brightness of images has been adjusted). (**B**) Representative images of nematodes colonised with the pLac biosensor strain. Top: uninduced, bottom: induced with 1mM IPTG. (**C**) Preliminary characterisation of the pLac biosensor. The first two columns refer to the negative and positive control, respectively. These are then followed by EcN_pLac_GFP_mCherry colonised worms, both uninduced and induced.(n *≥* 15 images, p-values: \*\*\**<*0.001).

**Figure 4:**
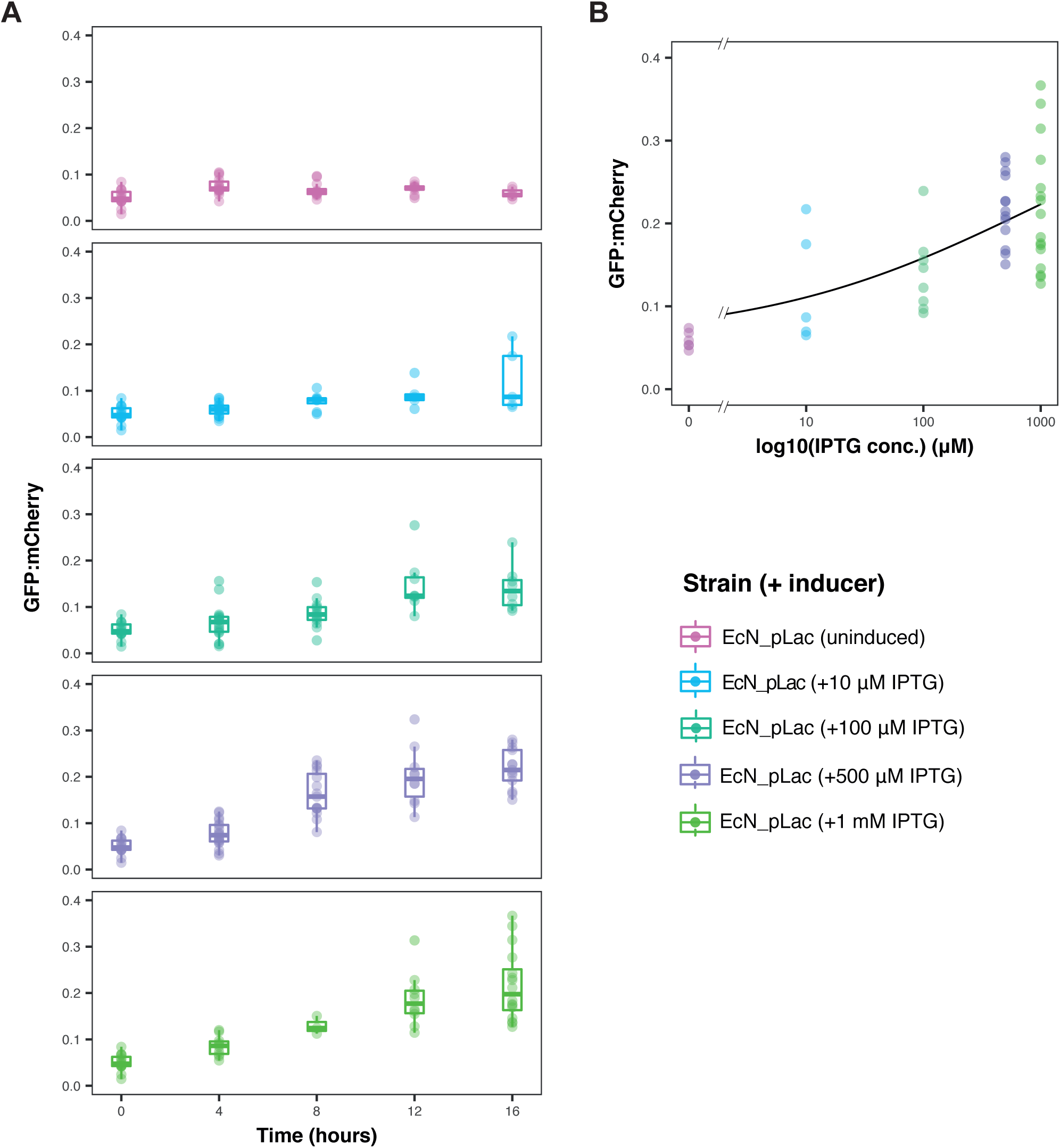
Full characterisation of the pLac biosensor *in vivo*. (**A**) Timecourse of GFP:mCherry ratios in individual 7 day old *C. elegans* worms, grown on EcN_pLac_GFP_mCherry sensor strain and transferred to inducer plates supplemented with varying IPTG concentrations. (n *≥* 4 images). (**B**) The GFP:mCherry ratios of the pLac biosensor, with various IPTG concentrations at the 16 hour timepoint, fit with a hill function (as in Figure 1).

The mean GFP:mCherry ratio of nematodes exposed to IPTG was 0.42±0.12, which was higher than the nematodes grown on the pLac strain without IPTG (0.14±0.04). In addition, the uninduced EcN_pLac_GFP_mCherry ratios were higher than those of the EcN_OG241_GFP_mCherry negative control strain demonstrating relatively high basal levels of GFP expression even when uninduced. This was also seen when comparing characterisation of the two strains *in vitro*, with EcN_OG241_GFP_mCherry showing lower ratios at both 20°C and 37°C (see Figures 1 and S4). This suggests the higher basal expression is caused by inherent properties of the plasmid system, rather than a factor of the environment present within the *C. elegans* digestive tract.

### Characterisation of the pLac (IPTG-inducible) biosensor *in vivo*

Our next goal was to demonstrate that the EcN_pLac_GFP_mCherry strain could detect and report on an environmental signal from within the *C. elegans* intestines in a quantitative manner. To ensure separation of the biosensor from the external environment, an induction assay was carried out whereby worms were grown for 7 days on nematode growth medium (NGM) agar plates spread with the pLac strain were transferred to plain NGM agar plates supplemented with varying concentrations of IPTG. The results are provided in Figure 4A. From these results it can be seen that the GFP:mCherry ratios increased in both a time and dose-dependent manner. This was not seen in the EcN_OG241_GFP_mCherry or EcN_OXB19_GFP_mCherry control strains (see Figure S7). Substantial increases in the GFP:mCherry ratios could be seen in as little as 4 hours, for as low as 10 *µ*M IPTG. Figure 4B shows the induction at the 16 hour time point as a function of the log of IPTG concentration; superimposed is a hill-function fit showing dose-dependent response with increasing levels of IPTG.

## Conclusions

Collectively, the results reported here indicate that the live *C. elegans* model can be used to characterise engineered biological systems and to design precise microbiota investigations. With minimal regulatory and time constraints, experiments can be carried out on a powerful yet simple and defined host-microbiota model system. This model has the potential to complement the underlying approach of the ‘design, build, test, learn cycle’ that is fundamental to synthetic biology.

The use of this model can provide valuable insights into general nematode biology. Currently, there is some debate within the nematode field as to whether *E. coli* become dormant within the digestive tracts of nematodes, after ingestion. As with the report by [29][29], the behaviour of the EcN dual-plasmid strains provide evidence that this is not the case. Data presented here suggests EcN is able to remain active even within the intestines of aged nematodes; with the EcN_pLac_GFP_mCherry strain capable of detecting and responding to environmental cues after a colonisation period of 7 days. Our results also suggest that, since GFP and mCherry require the presence of oxygen in order to fluoresce, the *C. elegans* intestinal environment is capable of transporting oxygen to some extent, even in older, fully colonised nematodes. Although this could hinder the ability of the model to predict sensor behaviour within the mouse or human intestines, the model can still be used to understand the performance of systems under normoxic conditions and could be developed further using nematodes that can survive in anoxic conditions [32]. Another finding is that some nematodes remained uncolonised using the protocol in Figure 2A. A possible explanation could be the aerobic state of the digestive tract, as densely colonised nematodes may not be capable of providing sufficient oxygen transport through their digestive tracts. This variation in colonisation may be explored further by investigating a range of colonisation periods. Future work will look at further exploration of the live *C. elegans* intestinal environment using engineered bacterial sensors.

The protocols described here have been used to characterise an IPTG-inducible biosensor. However, the system is readily adaptable to other possible metabolites or biomarkers of interest. In principle, this model could be used to detect inducers of interest introduced exogenously or derived from the host itself. IPTG is a molecule that is widely used as an inducer in synthetic genetic circuits. As the results of Figure 4 show, the pLac promoter can be induced while in the digestive tract of *C. elegans*.

Therefore, this promoter may be used in future studies to produce targeted expression within the digestive tract of the *C. elegans* nematode, in an analogous manner to how an aTc inducible system in *Bacteroides* was used in mice [14]. More generally, the *C. elegans* model will allow the exploration of host-microbe interactions and how bacterial strains compete within an *in vivo* environment. We believe that the methodology reported here will help expand our knowledge of the microbiome and allow for the reduction and replacement of current animal models used for testing *in vivo* synthetic biology approaches.

## Methods

### Flow cytometry analysis and plotting

Flow cytometry was performed on an Attune NxT Acoustic Focusing Cytometer, with Attune NxT Autosampler (Thermo Fisher Scientific, UK). 1 *µ*L of the appropriate strain culture was transferred into 200 *µ*L of sterile phosphate-buffered saline (PBS) in a shallow polystyrene 96-well plate. The Attune NxT Autosampler was used to record 10,000 events (for each individual sample) with 4 washes between samples. GFP was excited using the blue laser (488 nm) and detected using a 530/30 nm bandpass filter. mCherry was excited with the yellow laser (561 nm) and detected using a 620/15 nm bandpass filter. Additionally, a sample of 1:300 dilution of rainbow calibration particles in PBS (Spherotec, UK) was recorded allowing for the conversion of arbitrary units to MEF using Python scripts based on the FlowCal software [33].

Collected FCS data were analysed and plotted using custom Python and R scripts. Visualisation and curve fitting were performed in R, using the ‘ggplot2’ package and ‘nls’ fitting function. GFP induction and ratiometric increase data were fit using Hill functions

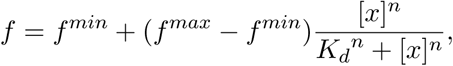

where *f* is the observed value (either fluorescence or ratio), *f* ^*min*^ is the minimum fitted value, *f* ^*max*^ is the maximum fitted value, [*x*] is the inducer concentration, *K*_*d*_ is the threshold sensitivity and *n* is the cooperativity (these parameters are illustrated in Figure S6). Dynamic range was calculated using the expression

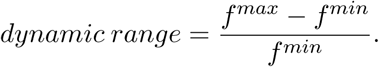

The linear range was calculated by taking the derivative of the Hill function over the length of the Hill fit

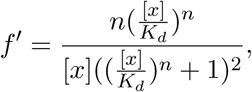

and then defined as the range of concentrations, over which the derivative of the Hill function was above 5% of the max value.

### *C. elegans* strains and handling

All *C. elegans* experiments were carried out in the wild-type lab strain N2 (provided by the Caenorhabditis Genetics Center, USA). Unless stated otherwise, worms were maintained and raised at 20°C on nematode growth medium (NGM) seeded with *E. coli* OP50, an auxotroph lab strain whose growth is limited on NGM.

Adult worms were maintained and passaged according to normal protocols. Briefly, this involved seeding NGM agar plates with 150 *µ*L of overnight *E. coli* OP50 culture and incubating for 48 hours at 20°C to create a bacterial lawn for food. Five to six L4-stage worms were picked and transferred to seeded NGM plates and incubated for 24 hours, allowing them to reach adult stage and begin laying eggs. After 24 hours, the adult worms were removed from each plate. The plates were then incubated for a further 48 hours to enable a large number of eggs to hatch, feed on the bacterial lawn and become fertilised with eggs.

After 48 hours, an ‘egg prep’ was carried out to isolate the eggs from the worms. This involved washing NGM plates with M9 media to collect all worms. Once settled, the M9 media was aspirated and the remaining worms were washed with 400 *µ*L of bleach:NaOH at a ratio of 7:8. This mixture was vortexed for 3-4 minutes (allowing release of the fertile eggs) and then neutralised with 13 mls of M9. The tube was then centrifuged until a firm pellet had formed and once again the M9 media was aspirated without disturbing the egg pellet. This wash was repeated twice more before the egg pellet was resuspended with 10 mls of M9 media and transferred into a sterile, empty petri dish for 24 hours incubation. This incubation allowed the eggs to hatch and arrested the worms at the first larval stage (L1). This provided an entirely synchronised nematode population. After 24 hours incubation, 400 L1s were seeded on the appropriate bacterial NGM plate for experiments or re-passaging. Before imaging, worms were passaged as sterile and non-reproductive adults with the drug fluorodeoxyuridine (FUdR). This involved picking worms at the L4 stage and transferring them to seeded NGM plates, supplemented with FUdR at a concentration of 20 *µ*M. This enabled the maintenance of a synchronously ageing population of worms.

Individual EcN-NGM plates were prepared in the same manner as with *E. coli* OP50 but with an overnight antibiotic culture of the respective EcN sensor strain or control instead. After preparing the worms as previously described, synchronised 400 L1 stage worms were transferred to NGM plates seeded with the respective EcN strain. These included EcN_OXB19_GFP_mCherry and EcN_OG241_GFP_mCherry as positive and negative controls, respectively. The inducible EcN_pLac_GFP_mCherry was used to investigate the sensor assay. After 48 hours of growth on the respective EcN-NGM plates, 45-50 L4 worms were picked and transferred to FUdR coated NGM plates seeded with the same EcN strain as before.

### EcN biosensor induction assays in *C. elegans*

As worms age, the peristaltic movements in their intestines decrease and they eventually become constipated while the bacteria proliferates in the gut [19]. Induction assays were carried out on 7 day old adult worms as they became colonised by the fast growing EcN. Initially plain NGM plates were prepared and seeded with the overnight culture of the relevant strain. These were then incubated at 37°C overnight. Approximately 50 sterile nematode eggs (gained from the egg prep mentioned above) were added to these seeded plates and incubated at 20°C for 2 days. Around 30 nematodes were then collected at random and transferred to EcN-NGM plates supplemented with the same strain, supplemented with FUdR to prevent egg maturation (as detailed above). These were then incubated for a further 5 days (to a total of 7 days), to allow for full colonisation of the majority of nematodes. It should be noted that antibiotics were not used in the EcN-NGM plates, instead it was assumed that the majority of the EcN bacteria would retain the dual plasmid system over this 7 day period. Approximately 25 worms were then picked from the respective EcN-NGM plates and transferred to either an unseeded plain NGM agar plate or an unseeded NGM agar plate containing the relevant IPTG concentration. Plates were then stored at 20°C for the duration of the assay. After induction, worms from either the control NGM plate or the assay plate were anaesthetised for imaging with 0.2% levamisole.

### *C. elegans* imaging

Anaesthetised worms were imaged with a Zeiss Axio Scope using GFP (excitation: 470nm; emission: 525nm) and mCherry (excitation: 560nm; emission: 630nm) filters. Exposure times were set at 500ms for each and laser intensities were kept constant. Images were acquired using the Zen software and analysed using the developed MATLAB pipeline. Further details on materials and methods can be found within the supplementary information.

## Acknowledgments and Funding

J.W.R. and T.O. were funded through the BBSRC LIDo doctoral training partnership. L.M.Q. was funded through the MRC. G.T. was supported by funding from the BBSRC and EPSRC. F.C. was supported through the Wellcome Trust/Royal Society (Grant No. 102531/Z/13/A) and the MRC (Grant No. MC-A654-5QC80). C.P.B. was supported through the Wellcome Trust (Grant No. 097319/Z/11/Z) and the European Research Council (ERC) under the European Union’s Horizon 2020 research and innovation programme (Grant No. 770835).

