## Supplementary Information for "Detecting changes in the *Caenorhabditis elegans* intestinal environment using an engineered bacterial biosensor"

### 1 Materials and Methods

#### Plasmids and Cloning Methods

Ratiometric sensor assay systems were designed with a constitutive mCherry reporter plasmid and a separate GFP reporter plasmid (plasmid maps can be seen in Figure S1). The GFP reporter plasmids were constructed from the promoterless OG241_GFP plasmid (Figure S1A), containing an upstream multiple cloning site, a dasher GFP reporter gene, a pUC high copy origin-of-replication and a kanamycin resistance cassette (Oxford Genetics, UK). As a positive control, the strong constitutive OXB19 promoter (Oxford Genetics, UK) was placed upstream of GFP to derive OXB19_GFP. In addition to these constructs, an isopropyl $\beta$-D-1-thiogalactopyranoside (IPTG) inducible GFP sensor (pLac_GFP) was created. Briefly, this was created by PCR cloning the region containing the LacI protein, tetR promoter and trc promoter from pKDL071 [1] and adding SpeI and EcoRI flanks. This region was then cloned into the MCS of OG241_GFP with the appropriate restriction enzymes. The p47_mCherry construct was created from pSEVA471, a 3120bp plasmid based on the pSEVA format [2]. This involved using PacI and SpeI to clone out the constitutively expressed mCherry fragment from pTn7-M-pEM7-mCherry and ligate into the empty multiple cloning site (MCS) of pSEVA471 to derive a low copy SC101 plasmid with streptomycin resistance and strong constitutive mCherry expression under the pEM7 promoter. EcN was then simultaneously transformed with p47_mCherry and the respective GFP plasmid via heat-shock methods to give the strains listed below.

Table S1: Bacterial strains used within this work.

| **Strain designation** | **Host** | **Plasmids** | **Source** |
|---|---|---|---|
| EcN_OG241_GFP_mCherry | *E. coli* Nissle 1917 | OG241_GFP_pUC_KanR, p47_M7_mCherry_SC101_StrpR | Oxford Genetics, UK |
| EcN_OXB19_GFP_mCherry | *E. coli* Nissle 1917 | OXB19_GFP_pUC_KanR, p47_M7_mCherry_SC101_StrpR | This work |
| EcN_pLac_GFP_mCherry | *E. coli* Nissle 1917 | pLac_GFP_pUC_KanR, p47_M7_mCherry_SC101_StrpR | This work |

### Media and Strains

Lysogeny broth (LB) media and agar were used during propagation and cloning of bacteria. Assays were carried out in LB or M9 minimal media supplemented with 1 mM thiamine hydrochloride, 0.4% glycerol, 0.2% casamino acids, 2 mM $MgSO_4$ and 0.1 mM $CaCl_2$. When antibiotic selection was applied, kanamycin was used at 25 $\mu$g/mL and streptomycin was used at 50 $\mu$g/mL. All DNA manipulations were performed in *E. coli* DH5$\alpha$ (NEB). All sequences were confirmed via Sanger sequencing (Source Bioscience, UK). All final induction assays were performed in *E. coli* Nissle 1917 (EcN) (Prof. Ian Henderson from the University of Birmingham, UK). Chemically competent EcN was made using standard protocols.

Briefly, this involved diluting an overnight EcN LB culture 1:100 into 50 ml of fresh LB media and grown at 37°C in a shaking incubator to an $OD_{600}$ of 0.25 to 0.3. The culture was then chilled on ice for 15 minutes and then centrifuged for 10 minutes at 5000 g and 4°C. The medium was then discarded and the cell pellet resuspended in 30 ml of cold 0.1 M $CaCl_2$ before being kept on ice for a further 30 minutes. This suspension was then centrifuged for 10 minutes at 5000 g and 4°C. The supernatant was once again removed and the cell pellet was resuspended in 3 ml of cold 0.1 M $CaCl_2$ solution with 15% glycerol. The final cell suspension was then aliquoted and flash frozen before being stored at -80°C.

### Induction Assays in Liquid Culture

All induction assay cultures were grown for 3 hours at 37°C (350 rpm shaking), before being induced and then transferred to 20°C (no shaking) or kept in the same conditions (37°C, 350 rpm shaking), at a volume of 500 $\mu$L in a sterile autoclaved 96-well deep square well (2.2 mL) polypropylene plate (BRAND®, Sigma-Aldrich, UK) sealed with Breathe-Easy sealing membranes (Sigma-Aldrich, UK). Induction was calculated through flow cytometry or plate-reader analysis. IPTG was used to induce the pLac_GFP plasmid. Concentration range induction assays were performed by diluting triplicate overnight cultures 1:500 into LB or supplemented M9 media, incubating for 3 hours (37°C) unless stated otherwise and inducing with the appropriate inducer concentration. The cultures were then allowed to incubate for 18 hours (at desired shaker speed and temperature) before flow cytometry analysis, as described below.

### Plate Reader Assays

Plate reader assays were performed in a Tecan Spark® plate reader (Tecan, Switzerland). Overnight cultures of uninduced bacteria were diluted 1:1000 into sterile LB media and incubated for 3 hours at 37°C. The cultures were then induced with the desired IPTG concentration and 200 $\mu$L of each culture transferred to a clear bottom 96-well plate (Greiner, Germany). Plates were then incubated at the desired shaking speed and temperature. Values for OD600, OD700, GFP (excitation: 488 nm, emission: 530 nm) and mCherry (excitation: 531 nm, emission: 620 nm) were recorded every 20 minutes over 20 hours.

### Biosensor Induction Quantification

The image analysis pipeline was developed using Matlab version R2016b and implemented on a 2015 MacBook Pro (8GB RAM). In brief, brightfield images were segmented to create a mask of the nematode bodies, this was then applied to the GFP and mCherry images. For images which contained segments of a worm already fully present within another image, this area was manually excluded (prior to placing within the automated pipeline) to ensure that every worm was only considered within a single image and to prevent oversampling. A threshold was also placed on object size, to ensure that small fragments of agar or eggs were not included in the worm mask. The colonised area of the mCherry intestine was then extracted and applied as a mask to the GFP image. GFP autofluorescence was accounted for by subtracting the background GFP signal of the worm body and uncolonised images discarded using an arbitrary mCherry threshold. Finally, the average pixel intensity of the GFP (green channel) and mCherry (red channel) for each image were collected in order to calculate the biosensor ratio. Where appropriate, the Mann-Whitney U test was used to demonstrate statistical significance. All plots were prepared using custom scripts based on the 'ggplot2' package in R.

### 2 Additional *in vitro* characterisation of EcN strains.

Increasing concentrations of IPTG can be used to achieve a maximum 25 fold increase in GFP fluorescence in EcN_pLac_GFP_mCherry with minimal impact on mCherry expression. Similar results were observed in induction with supplemented M9 media instead of LB (Figure S2). In a similar manner to the negative promoterless and positive constitutive controls, the uninduced EcN_pLac_GFP_mCherry strain demonstrated a lagged but steady level of mCherry expression over 15 hours (Figure S2C).

Further characterisation of the EcN_pLac_GFP_mCherry sensor in LB media, is presented within Figure S3.

Figure S4 shows characterisation of the negative (OG41) and positive (OXB19) control strains *in vitro*. From this figure it can be seen that while mCherry expression was identical in both the negative promoterless and positive constitutive control strains, the maturation of the red protein took far longer than the maturation of GFP. This delay in median mCherry fluorescence over the time-assay indicates that, while the production of mCherry may be constitutively and constantly driven by the promoter during cell growth, there is a delay in detection until the protein matures. In comparison, the fast maturation of GFP only results in a slight delay in detection during growth.

Even with the differences in maturation times, a distinctively higher GFP:mCherry ratio can be detected between the negative EcN_OG241_GFP_mCherry and the positive EcN_OXB19_GFP_mCherry control ($0.0125 \pm 0.0001$ compared to $0.471 \pm 0.004$ at 15 hours of growth) (Figure S4C). Due to the lag of maturation, the ratio of GFP:mCherry is particularly high during the logarithmic stages of growth (0 to 8 hours)(Figure S4C). In line with this, the difference in ratio between the strains is also apparent at the stationary phase of growth when the OD700 is larger than 1.5.

Figure S5 shows the parameters fitted during Hill Function modelling on a typical biosensor induction curve.

### 3 Additional *in vivo* characterisation of pLac biosensor.

Figure S6 shows the number of images which were discarded (deemed 'uncolonised' as below mCherry threshold) during image analysis, for each of the bacterial strains. It can be seen that only 17.23% of the total images were discarded. From this data no clear trends could be seen between percentage of 'colonised' images and either the strain, imaging timepoint or inducer concentration. This supports a hypothesis that the colonisation period may have a greater effect on the percentage of 'colonised' images.

Figure S7 shows characterisation of both the EcN_OG241_GFP_mCherry and EcN_OXB19_GFP_mCherry control strains, *in vivo* within the *C. elegans* digestive tract. From this characterisation it can be seen that the GFP:mCherry ratios for both control strains remain relatively constant, both in time and +/- the IPTG inducer.

Figure S8 contains the results of the original attempt to induce the pLac strain *in vivo*, after transferring to unseeded inducer plates. From the results it can be seen that there was no significant change in the GFP:mCherry ratio on transferring worms from seeded to unseeded plates, for a period of 8 hours without any inducer present. However, the addition of IPTG produced a significant increase in the GFP:mCherry ratio.

Figure S9 contains additional, higher magnification, images of a nematode colonised with the OXB19 control strain.

[2] Esteban Martinez-Garcia, Tomas Aparicio, Angel Goni-Moreno, Sofia Fraile, and Victor de-Lorenzo. SEVA 2.0: an update of the Standard European Vector Architecture for de-/re-construction of bacterial functionalities. *Nucleic Acids Research*, 43(Database issue):D1183–D1189, January 2015.

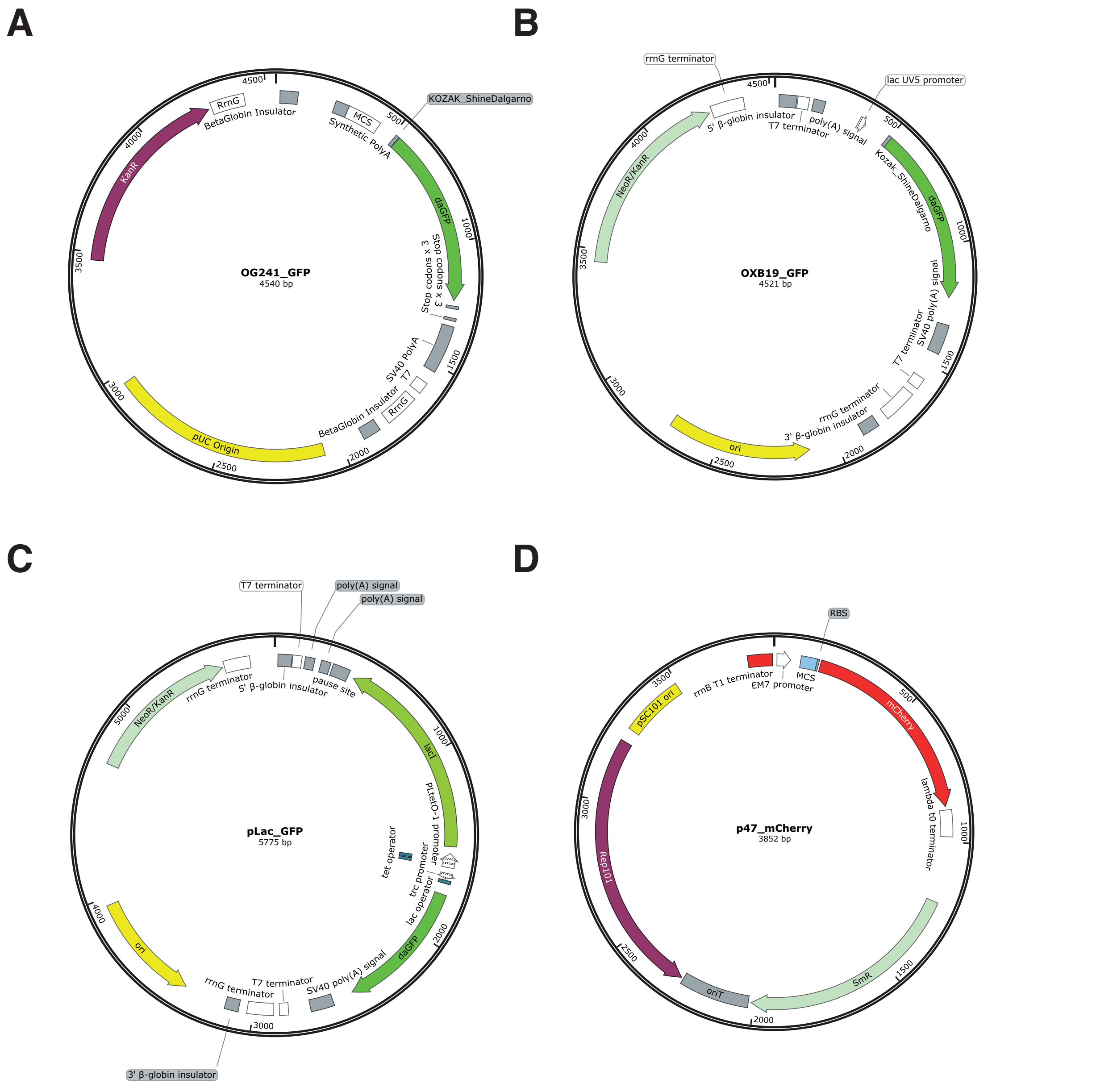


Figure S1: Maps of plasmids used within this work. (**A**) OG241, promoterless GFP. (**B**) OXB19, constitutive GFP. (**C**) pLac, IPTG inducible GFP. (**D**) p47_mCherry, constitutive mCherry.

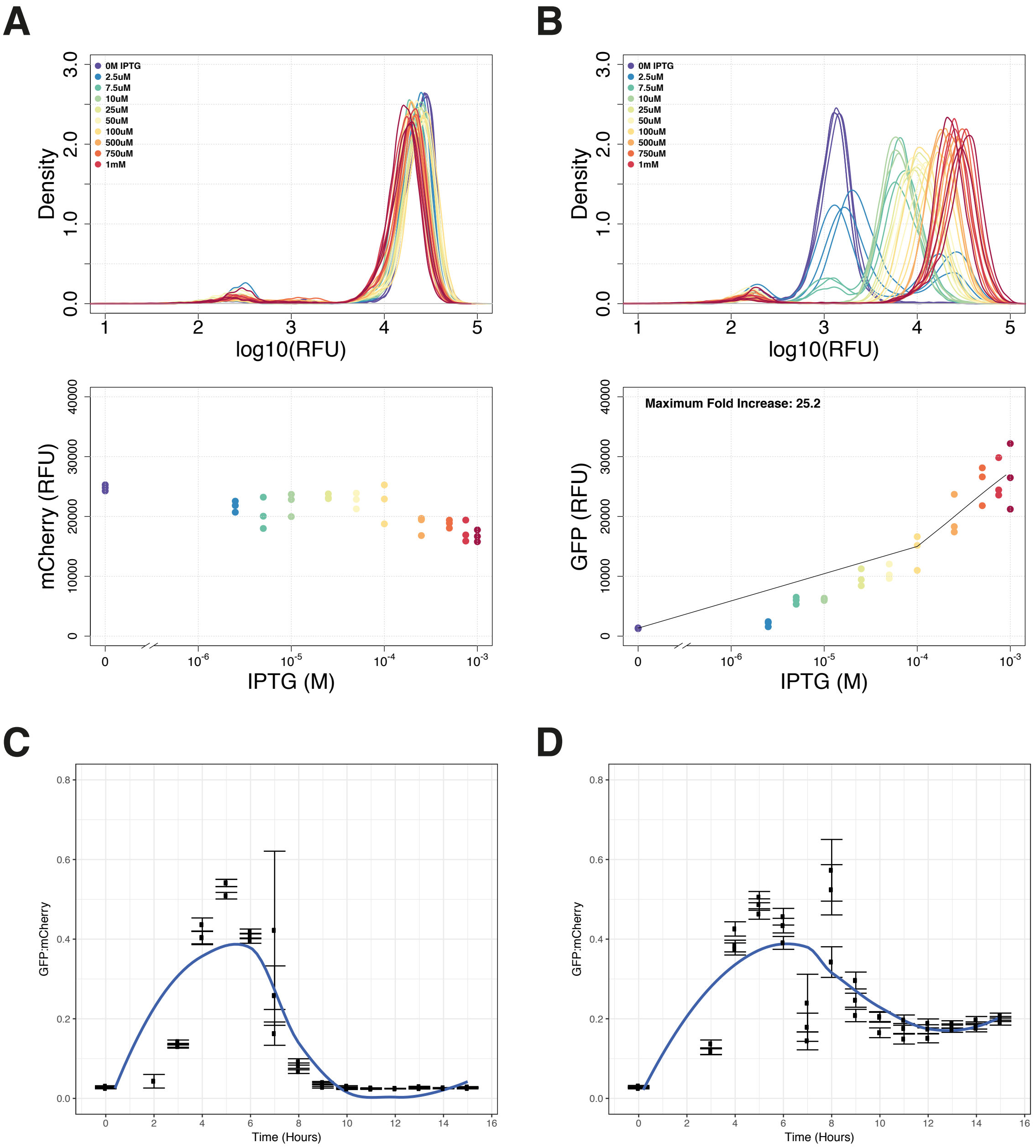


Figure S2: *In vitro* characterisation of EcN_pLac_GFP_mCherry, in M9 media. (**A**) mCherry expression after overnight IPTG induction. (**B**) GFP expression after overnight IPTG induction. (**C**) GFP:mCherry ratio in EcN_pLac_GFP_mCherry during growth, while uninduced. (**D**) GFP:mCherry ratio in EcN_pLac_GFP_mCherry during growth induced at 1mM IPTG at 7 hours in M9 media. Ratios were calculated using median fluorescence for each reporter from flow cytometry data with 10,000 events (n=3).

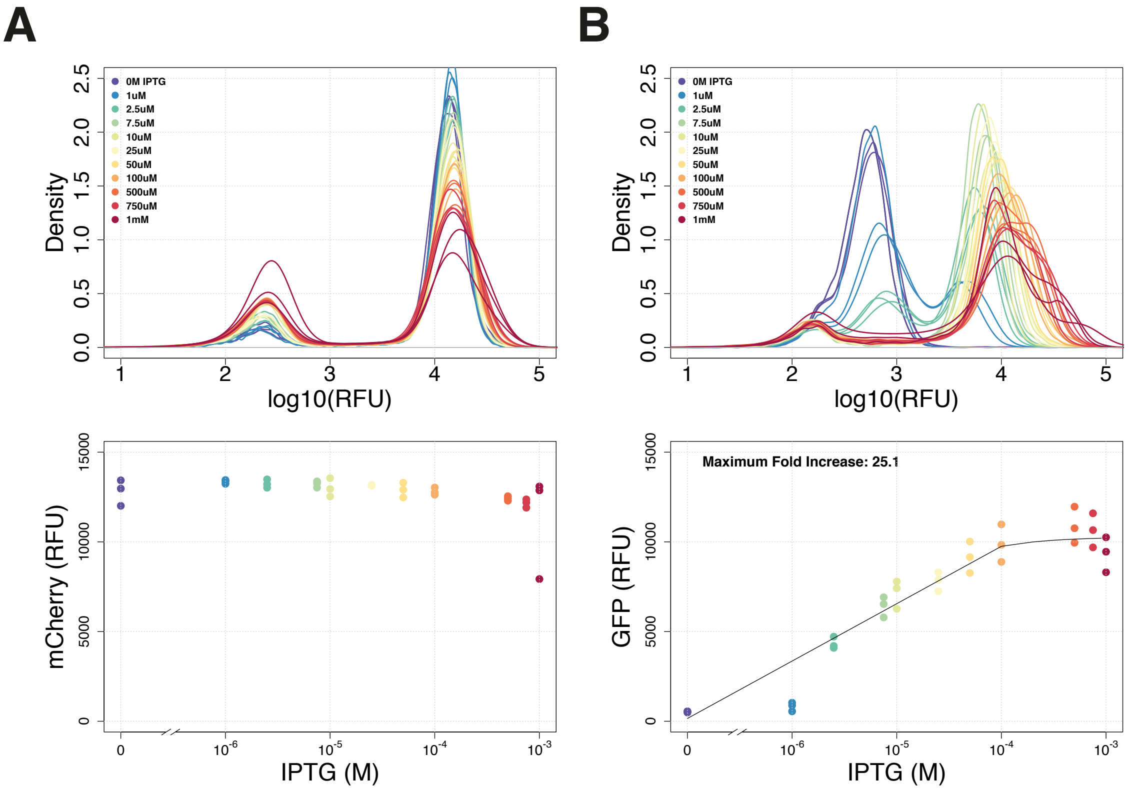


Figure S3: Further *in vitro* characterisation of EcN_pLac_GFP_mCherry, in LB media. (**A**) mCherry and (**B**) GFP expression after overnight induction with 1mM IPTG. Median fluorescence for each reporter was calculated from flow cytometry data with 10,000 events (n=3).

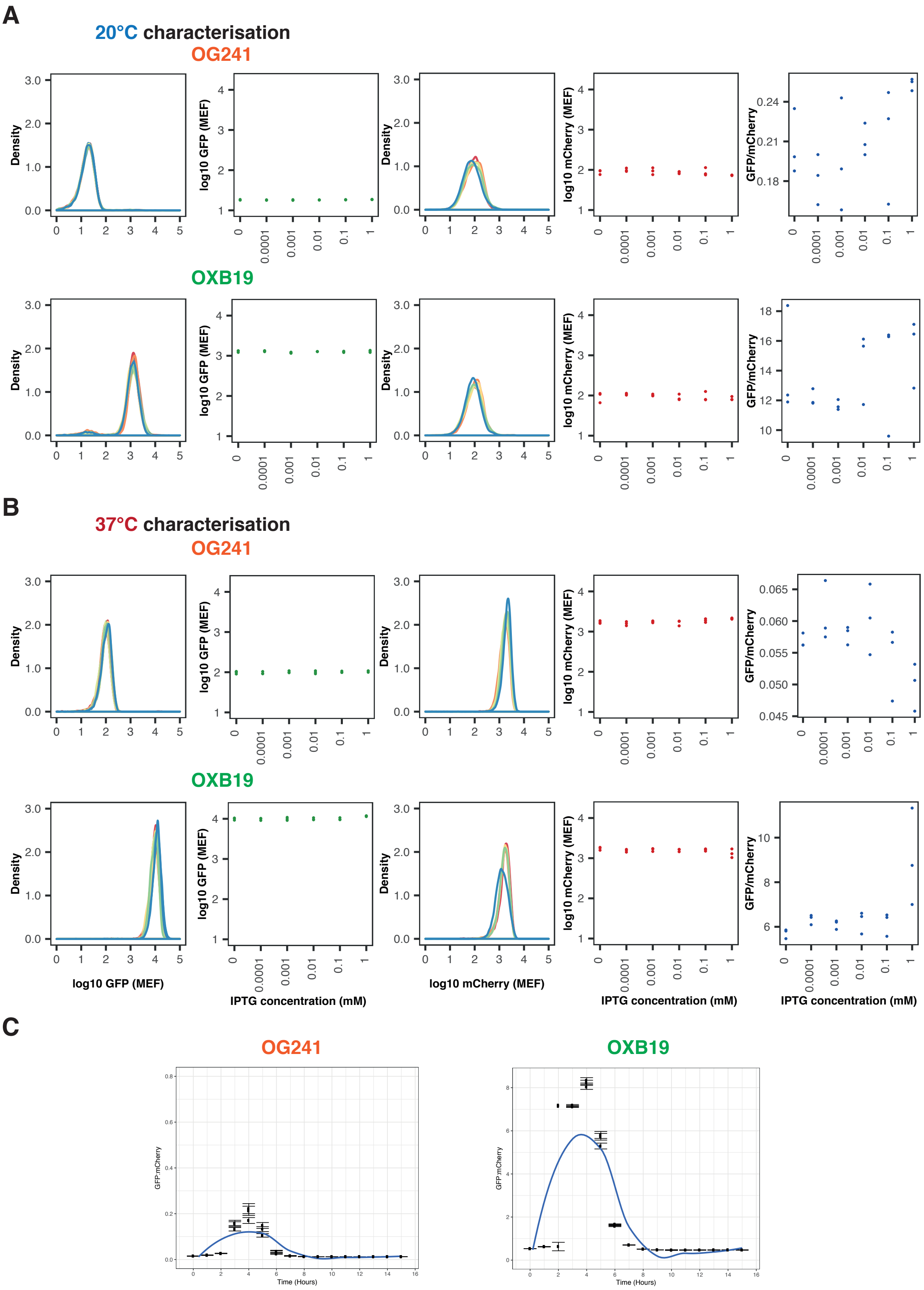


Figure S4: *In vitro* characterisation of the EcN control strains (EcN_OG241_GFP_mCherry and EcN_OXB19_GFP_mCherry). (**A**) 20°C, (**B**) 37°C, overnight induction assays. From left to right: density plot of GFP induction, median GFP fluorescence, density plot of mCherry fluorescence, median mCherry fluorescence and GFP:mCherry ratios over all IPTG inducer concentrations. (**C**) GFP:mCherry ratio timecourse, over 15 hours, for both strains at 37°C (not converted to MEF). The characterisation presented here was completed in LB media. Flow cytometry data with 10,000 events (n=3).

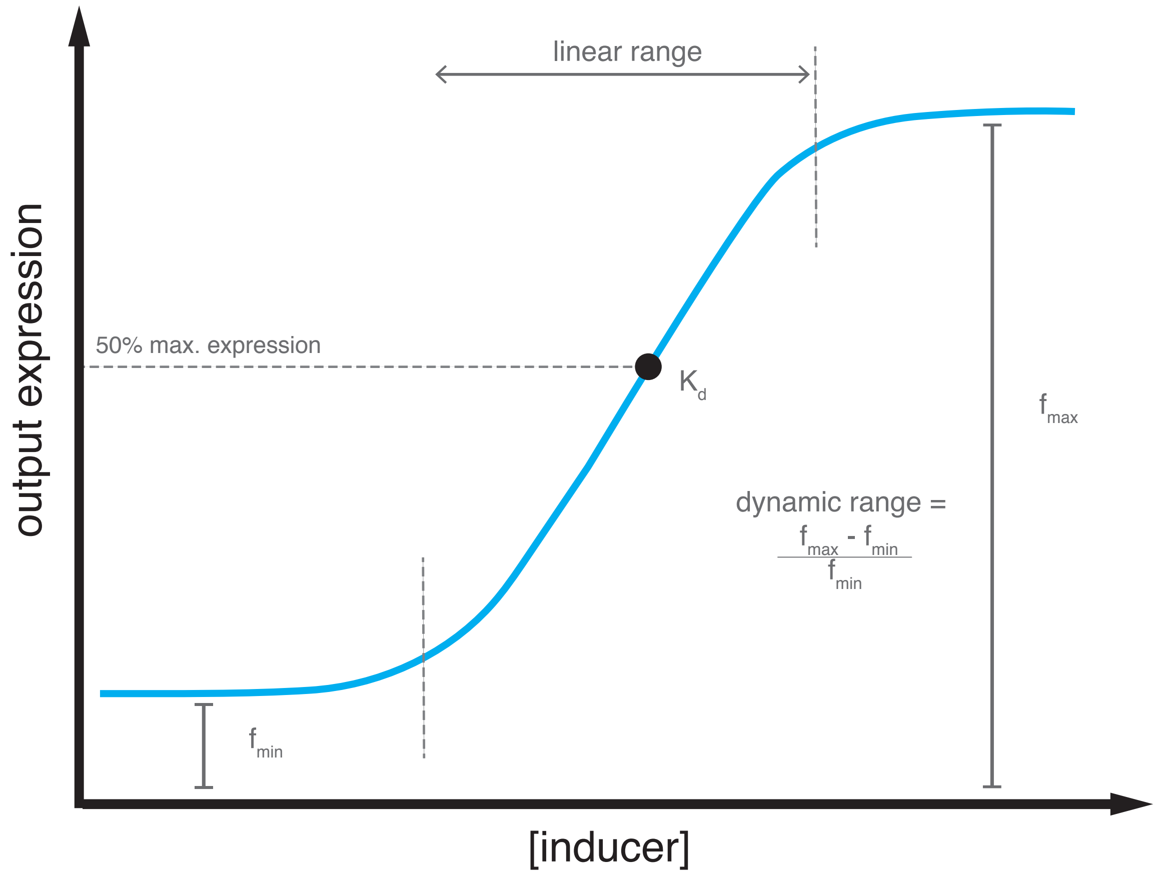


Figure S5: A typical biosensor induction response curve, highlighting the parameters estimated during Hill function fitting.

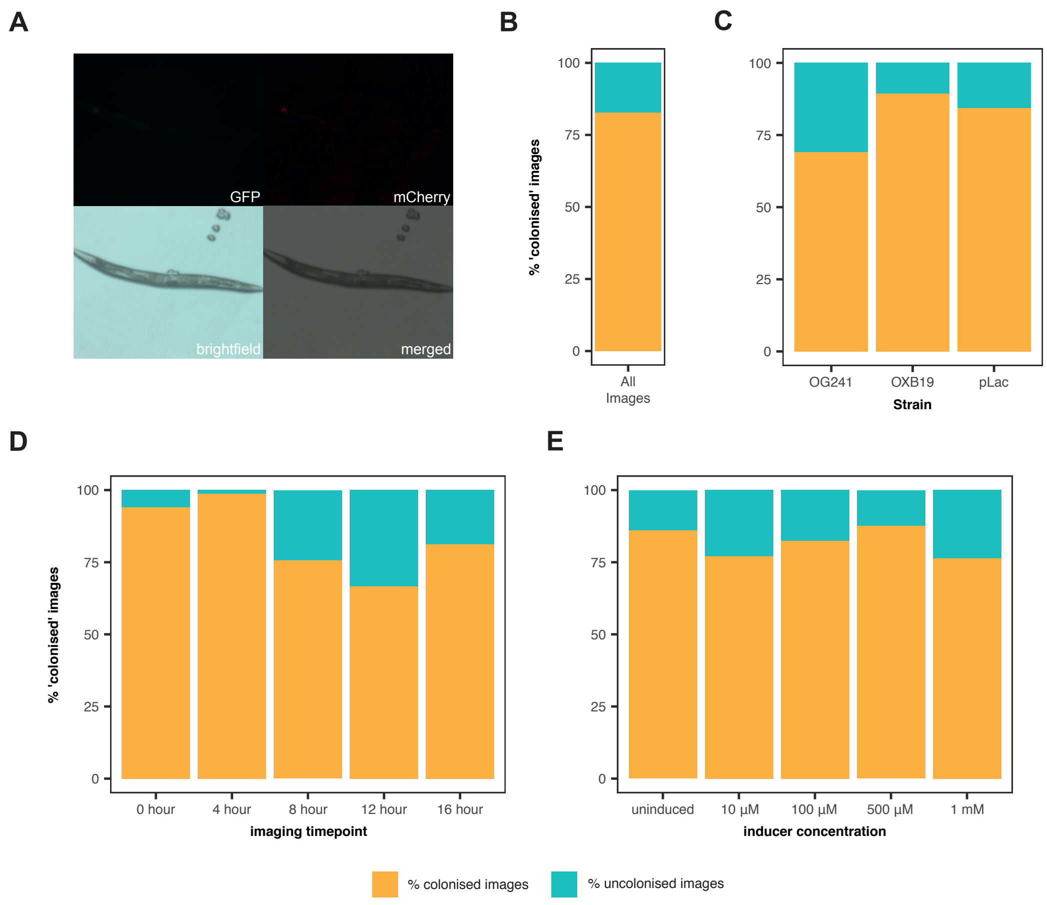


Figure S6: (**A**) Example image of a nematode which did not show mCherry fluorescence above the mCherry threshold and was therefore excluded during automated image analysis. Percentage of images which were kept ('colonised') or excluded ('uncolonised') during automated image analysis for each strain; for (**B**) all images, (**C**) images split by strain, (**D**) images split by timepoint, (**E**) images split by inducer concentration.

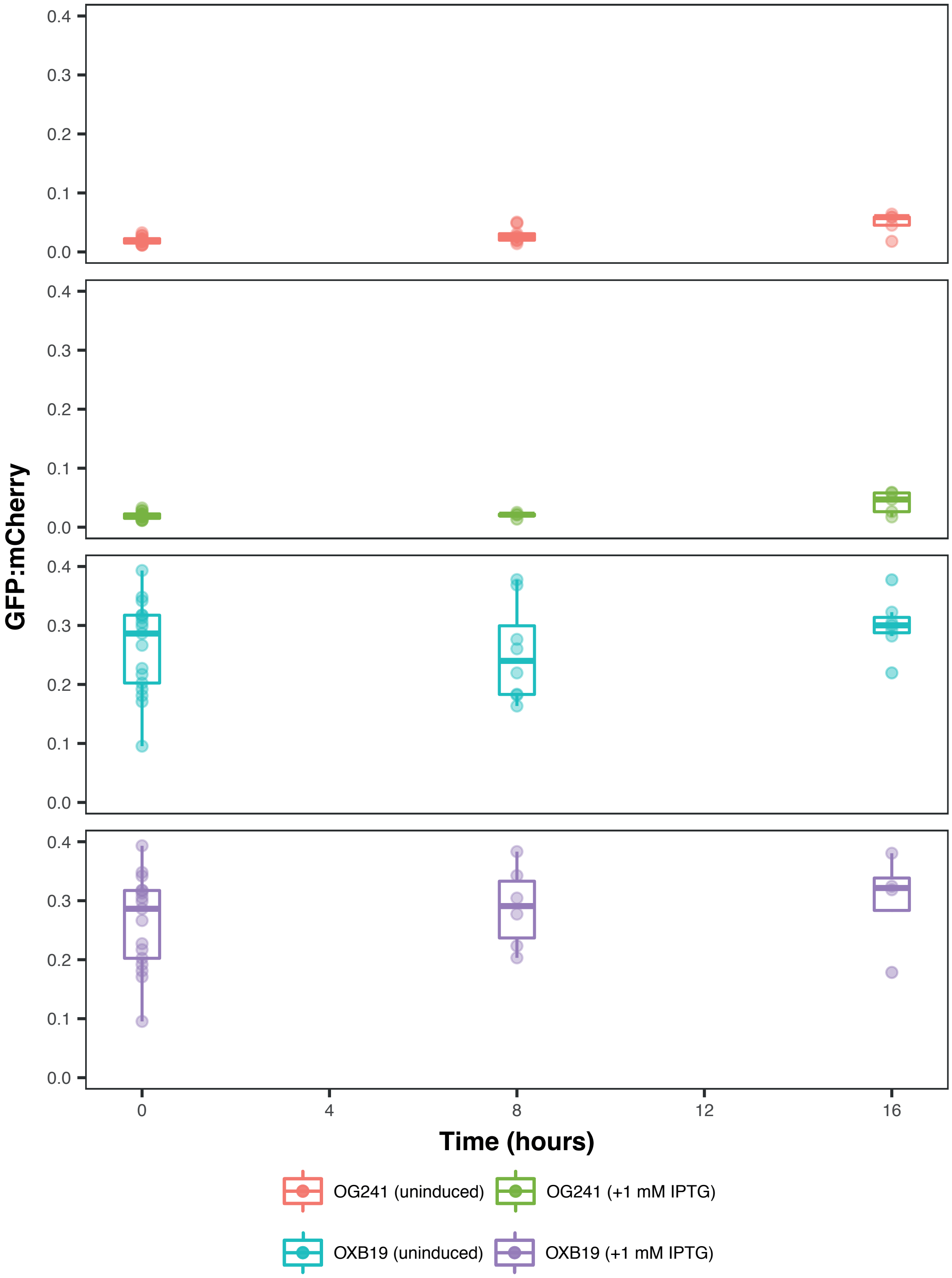


Figure S7: Characterisation of the EcN_OG241_GFP_mCherry (top) and EcN_OXB19_GFP_mCherry (bottom) control strains within the *C. elegans* digestive extract. Both with and without exposure to IPTG. (n $\geq$ 4 images).

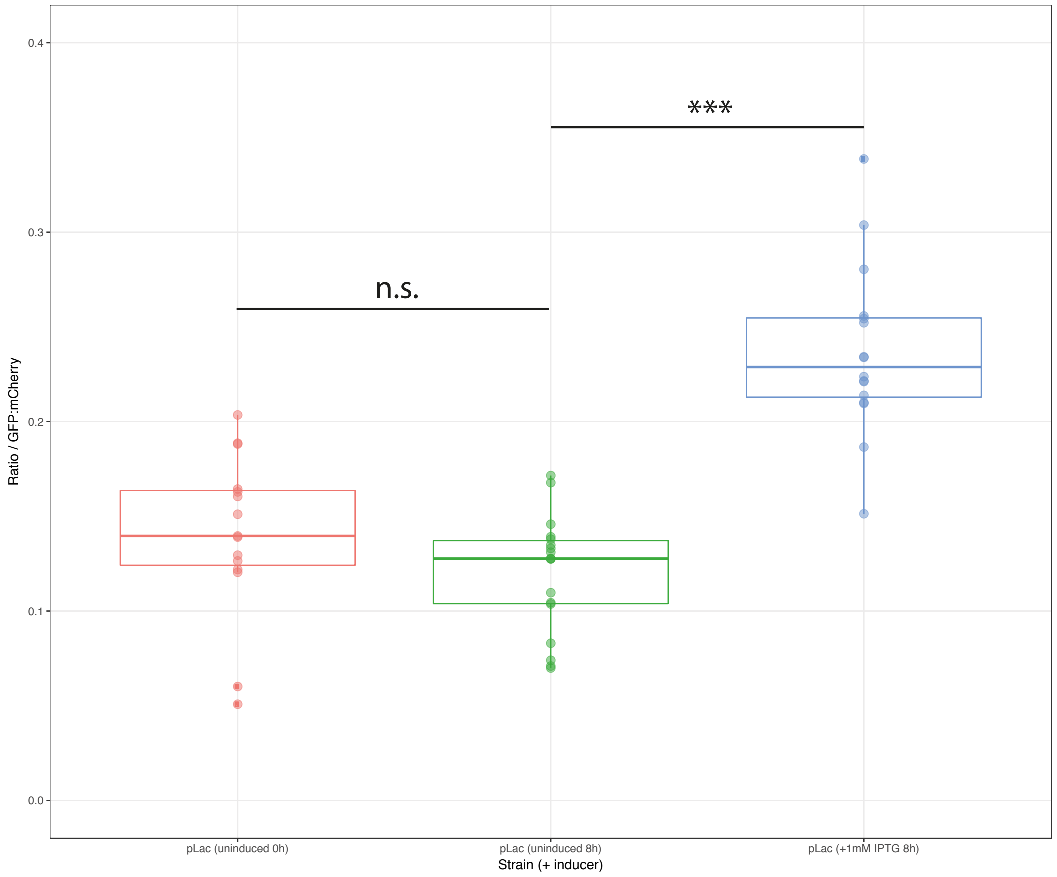


Figure S8: Preliminary characterisation of the pLac sensor within *C. elegans*. Images were collected at 0 hours and then 8 hours after transferring to inducer plates, supplemented with 0 or 1mM IPTG. The ratios of colonised nematodes are presented. ($n \geq 15$ images, p-values: n.s.>0.05, ***<0.001).

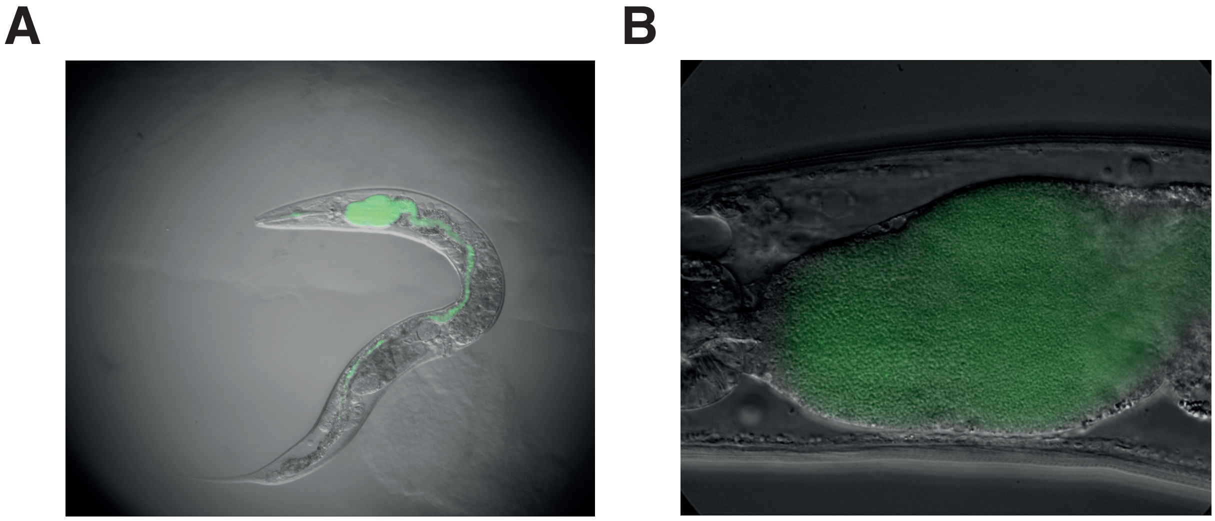


Figure S9: Images of colonised *C. elegans* nematode. Representative 7 day old *C. elegans* worm grown on EcN_OXB19_GFP, constitutively expressing GFP. GFP shows localisation of the strain from the pharynx onwards and throughout the intestines. (**A**) x120 magnification, (**B**) x200 magnification.
